# Co-dependent formation of the *Toxoplasma gondii* sub-pellicular microtubules and inner membrane skeleton

**DOI:** 10.1101/2024.05.25.595886

**Authors:** Klemens Engelberg, Ciara Bauwens, David J. P. Ferguson, Marc-Jan Gubbels

## Abstract

One of the defining features of apicomplexan parasites is their cytoskeleton composed of alveolar vesicles, known as the inner membrane complex (IMC) undergirded by intermediate-like filament network and an array of subpellicular microtubules (SPMTs). In *Toxoplasma gondii*, this specialized cytoskeleton is involved in all aspects of the disease-causing lytic cycle, and notably acting as a scaffold for parasite offspring in the internal budding process. Despite advances in our understanding of the architecture and molecular composition, insights pertaining to the coordinated assembly of the scaffold are still largely elusive. Here, *T. gondii* tachyzoites were dissected by advanced, iterative expansion microscopy (pan-ExM) revealing new insights into the very early sequential formation steps of the tubulin scaffold. A comparative study of the related parasite *Sarcocystis neurona* revealed that different MT bundling organizations of the nascent SPMTs correlate with the number of central and basal alveolar vesicles.

In absence of a so far identified MT nucleation mechanism, we genetically dissected *T. gondii* γ-tubulin and γ-tubulin complex protein 4 (GCP4). While γ-tubulin depletion abolished the formation of the tubulin scaffold, a set of MTs still formed that suggests SPMTs are nucleated at the outer core of the centrosome. Depletion of GCP4 interfered with the correct assembly of SPMTs into the forming daughter buds, further indicating that the parasite utilizes the γ-tubulin complex in tubulin scaffold formation.

## Introduction

The phylum Apicomplexa is largely comprised of obligate intracellular parasites with a defining apical constellation of secretory organelles and cytoskeletal structures uniquely evolved to facilitate invasion of another host cell. The assembly of these apical elements is orchestrated by the centrosome and unfolds through a budding process wherein the cortical cytoskeleton serves as a scaffold for *de novo* assembly of secretory organelles [1]. Although there is significant customization of the parasite size and budding mode to each parasitic niche, the assembly of the cortical cytoskeleton is the principal driver of budding and thus conserved across the phylum [1–3]. The cortical cytoskeleton is not only important for cell division (budding) but is also responsible for gliding motility required for successful host cell invasion [4]. Hence, it is pivotal for the success of apicomplexan parasites.

The cortical cytoskeleton is categorized as an epiplastin-based membrane skeleton, which is principally different from the actin-spectrin-based cytoskeleton present in mammalian cells [5]. The three main elements of the parasite’s cytoskeleton are a set of flattened alveolar vesicles (alveoli) upheld by a set of intermediate filament-like proteins (named alveolins in general, but in Apicomplexa known as alveolin-domain containing inner membrane complex (IMC) proteins) [6–9] and sub-pellicular microtubules (SPMTs) anchored by the apical polar ring (APR) at the apical end of the parasite [10, 11]. The number of SPMTs and alveolar plates is highly variable across parasite species and developmental stages with the same species [2]. The apical end of the cytoskeleton is characterized by the conoid, with various elaboration degrees in different parasites, composed of unconventional tubulin-fibers and two polar rings [12].

The tachyzoite stage of *Toxoplasma gondii* is an established model apicomplexan in which many of the cytoskeleton and cell division details have been unraveled [1, 4, 6]. In the asexual stages of *T. gondii* and related coccidian parasites like *Neospora caninum* and *Sarcocystis neurona*, the number of SPMTs is 22, while the number of alveolar plates varies [2, 13–15]. The IMC (alveoli together with alveolin-domain containing IMC proteins) rests on the SPMTs that radiate from the APR [10] and are evenly spaced in the mature parasite.

The *T. gondii* tachyzoite divides by arguably the least complex mode of apicomplexan cell division, endodyogeny, forming two new daughter parasites per one round of division within the cytosol of the mother cell [1, 6, 16]. Each daughter buds from a bipartite centrosome that is divided in the inner core, which controls the nuclear cycle, and the outer core, controlling bud assembly [17, 18]. Bud formation is dependent on two proteins homologous to algal striated fiber assemblin (SFA) that form a fiber positioning and anchoring the bud scaffold to the centrosome [16, 18–20]. Bud formation is initiated by scaffolding and signaling proteins, which are initially laid out in a 5-fold symmetry [21–26] (**Fig 1A**). The daughter buds grow first bi-directional to later adapt a strictly apical to basal growth trajectory [16, 21, 27]. All three elements of the cortical cytoskeleton are intertwined and appear to assemble in parallel rather than sequentially [21, 28, 29]. However, the understanding of early bud formation and especially how the assembly of the three elements is coordinated, is limited. Concomitant with the APR and SPMTs, the conoid is formed, consisting of 14 unconventional tubulin fibers [30] and two intra-conoidal microtubules (ICMTs) [31]. With progress of division, the dividing nucleus is encapsulated by the forming daughter buds (mitosis and cytokinesis progress in parallel) [32]. At the basal end of the forming daughter buds lies the functional equivalent of the contractile cytokinetic ring, the basal complex (BC), which is responsible for yoke the (+)-ends of the SPMTs and later on for tapering and constricting the nascent daughters [21, 33–37]. Division concludes with disassembly of the mother’s IMC followed by the daughter parasites emerging whilst depositing their mother’s plasma membrane on their IMC [38, 39].

**Fig 1.**
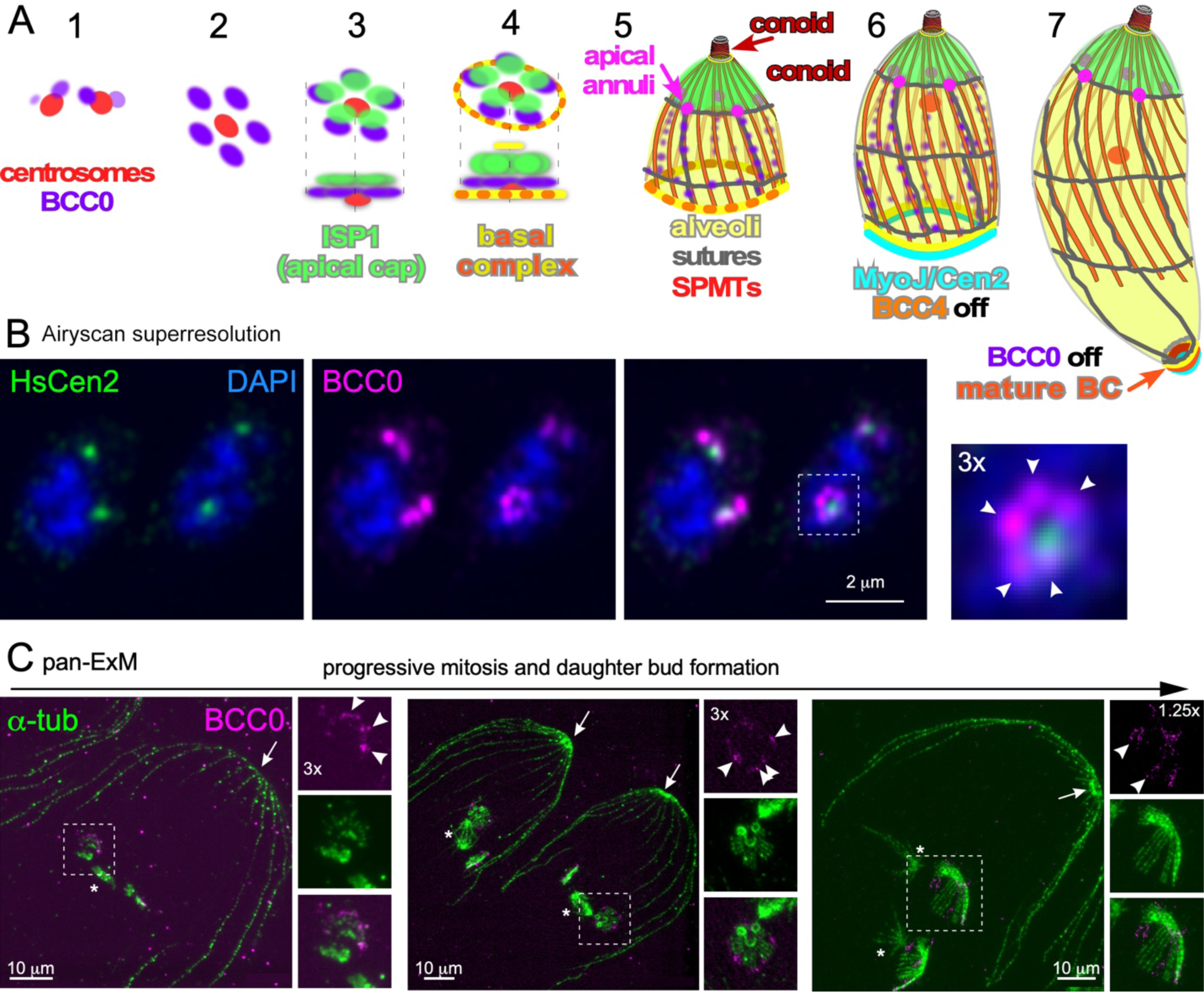
Early daughter cytoskeleton assembly displays five-fold symmetry, alternating between nSPMT rafts and the foundation of longitudinal alveolar sutures. **A.** Schematic representation of *T. gondii* cytoskeleton assembly. Stage 1-7 represent advancing steps in daughter cytoskeleton scaffold assembly. Modified from [21]. **B.** Original observation of BCC0 bud initiation as five distinct puncta, with five-fold symmetry (puncta marked by arrowheads). **C.** Pan-ExM of three different parasites progressing through mitosis and early bud assembly. Asterisks mark mitotic spindle; arrows mark the apical end of the mother’s cytoskeleton (conoid); arrowheads mark the BCC0-5xV5 signals deposited before nSPMT assembly in five foci around the centrioles, which subsequently accumulates between the nSPMTs rafts

How the SPMTs are nucleated is still not known in detail, but the APR has been proposed as an unusual microtubule organizing center (MTOC) in which the (−)-ends of SPMTs are inserted between blunt projections to form a cogwheel-like pattern [10, 40–42]. This assignment is supported by its close apposition to the SPMTs and conoid as well as the presence of a centrosomal SAS6-like (SAS6-L) protein in the APR (the centrosome harbors its own SAS6) [43]. This insight together with several other suggestions and the key role for an SFA fiber indicates that daughter budding either has shared ancestry, or at least co-opted elements and mechanisms, shared with cilium formation in eukaryotic cells [44]. Several proteins have been identified that facilitate stability of the APR (Kinesin A, APR1, RNG2, AC9 and 10), but none affects bud initiation [22, 42, 45, 46]. Stability of SPMTs is modulated by several previously identified proteins, but similar to the APR, for none a detailed role in bud initiation has been established [28, 41, 47–50].

γ-Tubulin and its associated complex (γ-TuC) is a known nucleator of MTs [51], and γ-TuC key components (γ-tubulin, γ-tubulin complex proteins (GCP) 2, 3, and 4) have been identified in the *T. gondii* genome [52]. γ-Tubulin has previously been localized to the outer core of the centrosome [18]. Its molecular function, however, has not yet been established. Given the close proximity between the centrosomal outer core and the forming daughter bud, we hypothesized that γ-TuC may be functionally involved in the assembly of the tubulin scaffold.

We have recently adopted an advanced iterative expansion microscopy (ExM) technique, called pan-ExM [53]. Pan-ExM relies on two consecutive rounds of sample expansion, which allowed us to physically increase parasite size by up to a factor 13. Using this new technology, we have revealed the organization of the 22 nascent SPMTs (nSPMTs) in five distinct “rafts” around the APR at bud initiation. We compared this organization to *S. neurona* merozoites, which put down 11 pairs of two SPMTs, an organization that mirrors twice the number of alveolar sheets [14]. Our data further show that not all nSPMTs form at the same time, but rather are sequentially assembled on the APR, indicating a distinct area of nucleation. When we examined the localization of γ-tubulin by pan-ExM we observed it at the spindle poles, the centrioles, and, unexpectedly, also at the nascent APR (nAPR), specifically at the onset of division. Rapid depletion of γ-tubulin abolishes the formation of the tubulin scaffold and results in paired SPMTs emanating from centrioles. Thus, γ-tubulin localization and function highlight it as a pivotal component of daughter cell formation. Functional dissection of the SFA component SFA2 and γ-tubulin complex protein 4 (GCP4) further allowed us to gain insights into the assembly hierarchy of the daughter scaffold.

Overall, the combination of advanced imaging and genetic dissection provides deep new insights in how SPMT and alveolar architecture are coordinated to form the template for daughter bud formation.

## Results

### Five-fold symmetry of the *T. gondii* tachyzoite early daughter scaffold

We recently reported that the *T. gondii* daughter-specific protein BCC0 assembles in a distinct five-fold symmetry around the duplicated centrosomes early in division. Upon progression of budding BCC0 is deposited in the longitudinal IMC sutures of the daughter bud (**Fig 1A,B**) [21]. To refine our understanding of these very early steps in daughter cell assembly, we adopted pan-expansion microscopy (pan-ExM) [53–55], which in our hands results in an approximately 13-fold increase of sample size. Co-staining of α-tubulin and BCC0 through early daughter budding using pan-ExM revealed that BCC0 localizes in between distinct SPMT ‘rafts’, composed of four individual SPMTs, that are formed around the centrioles (**Fig 1C**). Temporally, templating of the SPMTs starts while the metaphase mitotic spindle is present. When the spindle poles separate during anaphase, the organization of SPMTs in rafts of four nSPMTs, as described recently by U-ExM [56], is still visible, but the nascent scaffold transitions from a flat plane into a cone-shape conformation. BCC0 remained visible between SPMT rafts, which is suggestive of coordination between BCC0 and SPMT formation (**Fig 1C**).

Next, we delved into the earliest steps in daughter scaffold formation, as pan-ExM permits viewing these early steps without visual perturbation from the maternal cortical cytoskeleton. Upon staining the MTs with α-tubulin, pan-ExM clearly resolves the two centrioles in the *T. gondii* centrosome, which are marked with centrosome marker HsCen2 forming a fiber between the centrioles [21] (**Fig 2A**). Also, α-tubulin is present in the mitotic spindle and highlights the nSPMTs. In line with recent reports [56, 57], the 22 nSPMTs are organized in five distinct rafts of propeller-like symmetry, consisting of four rafts (each four nSPMTs) plus one raft with six nSPMTs (**Fig 2A** [middle panel]) or five sheets of four SPMTs plus a sheet of two separate SPMTs. We noted that in the earliest stage shown in **Fig 1A**, the nSPMTs are very few in number, and very short. Subsequently, we looked for earlier steps in nSPMT formation and found several cases of three rafts (each with four nSPMTs) + two individual nSPMTs and four rafts (each with four nSPMT) assembled around the forming conoid (**Fig 2B**, arrowhead). At this stage, the nascent scaffold is reminiscent of a horseshoe and on one side in close proximity to one of the centrioles (**Fig 2B**). This horseshoe was recently assigned to be the early conoid [56]. We frequently observed a pair of two short MTs, overlaying with a centriole (arrows in **Fig 2B** left and center panels), and others apparently being added to one side of the forming scaffold (**Fig 2B**, yellow arrowhead). Taken together, we interpret these data into a model wherein the nSPTMs grow in pairs of two and are subsequently organized into rafts of four nSPMTs, whereas two additional nSPMTs are either on their own or added to another nSPMT raft to make a total of six. Germane to the early organization of nSPMTs, this suggests a defined area of nucleation, and a symmetrical coordination with BCC0 laying down the IMC. This subsequently outlines the pattern for the growing cytoskeleton, i.e. the position of the apical annuli and alveolar sheets (e.g. **Fig 1A**).

**Fig 2.**
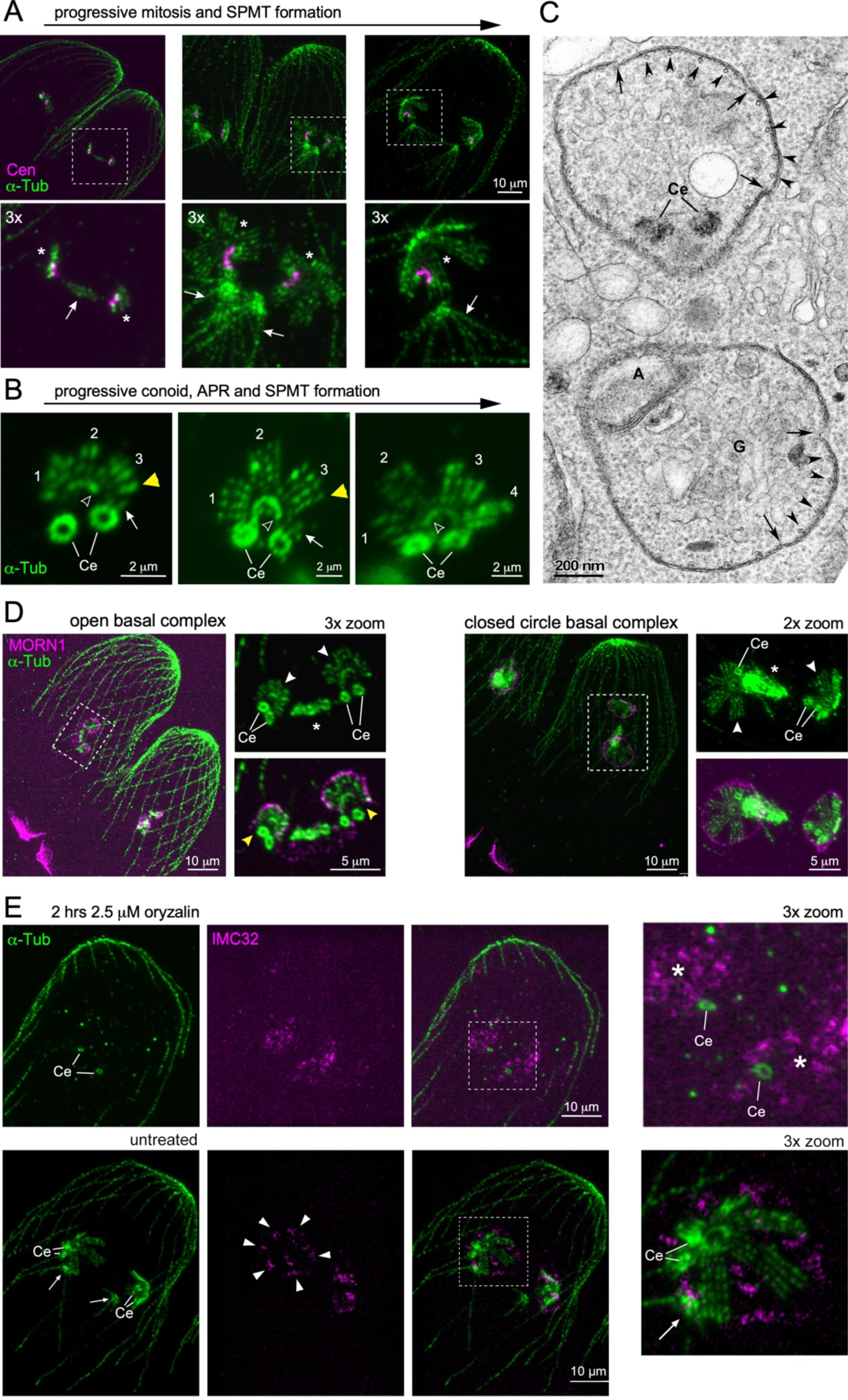
The nSPMTs display a five-fold symmetrical organization early in the division, in connection with the IMC and the basal complex. **A.** Pan-ExM of early budding parasites co-stained with *Hs*Centrin2 (Cen) and α-tubulin (MAb 12G10) reveals that the centrioles are connected by a centrin fiber. SPMT formation starts during mitotic anaphase (left panel; mitotic spindle between the centrosomes). Five-fold symmetrical rafts of nSPMTs assemble during telophase (middle and right panels), which transitions from a flat organization (middle panel) into a domed orientation (right panel). Arrows: mitotic spindles; asterisks: nSPMT rafts. **B.** Pan-ExM snapshots of sequential stages of conoid (open arrowheads) and nSPMT formation. nSPMT rafts are numbered. Arrows mark pairs of short MTs that are being added to the growing scaffold. Note that the ICMT pair is visible within the conoid in the middle panel. Ce: centrioles. **C.** Transmission electron microscopy of early daughter budding. Cross section through daughter buds wherein nSPMT rafts are visible with IMC membrane sheets on top. Arrowheads mark individual nSPMTs; arrows mark breaks in the membrane sheets of the nascent IMC alveoli. A: apicoplast, Ce: centriole; G: Golgi apparatus. **D.** Pan-ExM of the MT cytoskeleton co-stained with MORN1, highlighting how the nascent basal complex interfaces with the early scaffold (nSPTMs). Ce: centrioles; asterisks mark mitotic spindle; white arrowhead mark the nSPMT rafts; yellow arrowheads mark the discontinuity of the nascent basal complex assembled close to the (+)-end of nSPMTs. **E.** Pan-ExM reveals the differential susceptibility of microtubules to Oryzalin treatment. As previously observed [60, 61] division is not halted by this treatment, but cytoskeleton organization is substantially altered. After oryzalin treatment, IMC32-5xV5 forms plaques (asterisks) around (single) centrioles, contrary to its localization in discrete foci (arrowheads) between nSPMT rafts in untreated controls. Ce: centrioles. arrows: mitotic spindle;

### SPMTs assembly pattern provides foundation for alveoli plate architecture

The five-fold symmetrical nSPMT organization in **Fig 1C** correlates with the five distinct foci, previously detected for BCC0. To directly visualize the membrane of the IMC we performed transmission electron microscopy (TEM). Indeed, we observed four nSPMTs undergirding membrane sheets with gaps between them (**Fig 2C**). From this we inferred that BCC0 localizes to the gaps as it becomes imbedded in the sutures between the 5-6 alveolar plates of the two rows of central and one row of basal alveoli [21]. In addition, the TEM showed a centriole pair in close apposition to one side of the early daughter bud, mirroring our pan-ExM observations in **Fig 2B**.

Another early and crucial component of daughter buds is the scaffolding protein MORN1, which assembles the BC at the basal end of the cytoskeleton scaffold [21, 33, 58, 59]. During the early stages while the nSPMT are still forming, we observed MORN1 as a horseshoe-shaped open circle around the (+)-ends of the nSPMTs (**Fig 2D**, left panels). Once all nSPMTs are formed, MORN1 transitioned into a closed circle configuration of the mature BC (**Fig 2D**, right panels). However, at this point the nSPMTs are still organized in rafts, indicating the even spreading of SPMTs around the nascent nAPR must occur at a later stage and is independent of BC circle closure.

To further probe the correlation between nSPMTs and the alveolar architecture, we chemically perturbed SPMT formation using the MT destabilizer oryzalin [60, 61]. We then assessed IMC formation using the early marker IMC32, a membrane-anchored protein essential for daughter bud initiation displaying the same early symmetry as BCC0 [21, 23]. In parasites treated for 2 hrs with oryzalin, we observed single centrioles that are far separated from each other, indicating that, surprisingly, centrosome splitting progressed normally despite the lack of centriole duplication (**Fig 2E**). Centrosomal splitting indicated that budding had been initiated, although the spindle and nSPMTs were absent. IMC32 however, still accumulated around the single centrioles but did not display the typical symmetry, seen in the untreated controls (**Fig 2E**) [23]. Collectively, these observations support a model wherein the nSPMT organization defines the symmetry of the alveolar membrane organization.

### Comparative biology of nSPMTs and alveolar architecture

To further cement the connection between nSPMTs and the IMC alveolar sheet architecture, we turned to the closely related coccidian parasite *Sarcocystis neurona*. *S. neurona* is the causative agent of equine protozoal myoencephalitis [62]. Unlike *T. gondii* tachyzoites, *S. neurona* merozoites replicate by endopolygeny, where the parasite undergoes several rounds of S-phase and mitosis without completing karyokinesis, followed by a final round of S- and M-phase coupled to daughter budding, resulting in 64 emerging merozoites (**Fig 3A**) [1]. In relation to the question at hand, *S. neurona* merozoites, like *T. gondii* tachyzoites, harbor 22 SPMTs, but have twice as many alveolar sheets: 11 total (visualized in **Fig 3B** using the IMC suture marker TgIMC15-YFP [63]) vs 5-6 for *T. gondii* tachyzoites [15]). Hence, we anticipated the nSPMT to reflect the increased number of alveoli. This is indeed what we observed early in daughter cytoskeleton assembly, where nSPMTs formed 11 pairs, each consisting out of two nSPMTs. Like in *T. gondii*, the BC marker MORN1 is located close to the (+)-end of SPMTs during development and highlights the basal end of the forming bud. The observed MT ‘pairing’ was maintained through the mid-budding stage until budding is completed and the merozoites matured (**Fig 3C**). Interestingly however, the SPMTs in mature merozoites were alternating between a SPMT of about 2/3 the length of the cell with the other SPMT that extended all the way to the end of the merozoite, connecting to the basal complex (right panel, **Fig 3C**). This contrasts with *T. gondii* tachyzoites, where none of the SPMTs reach to the very basal end and only extent to 60% of the length of the mature tachyzoite, although some variation in SPMTs length is seen here as well [14].

**Fig 3.**
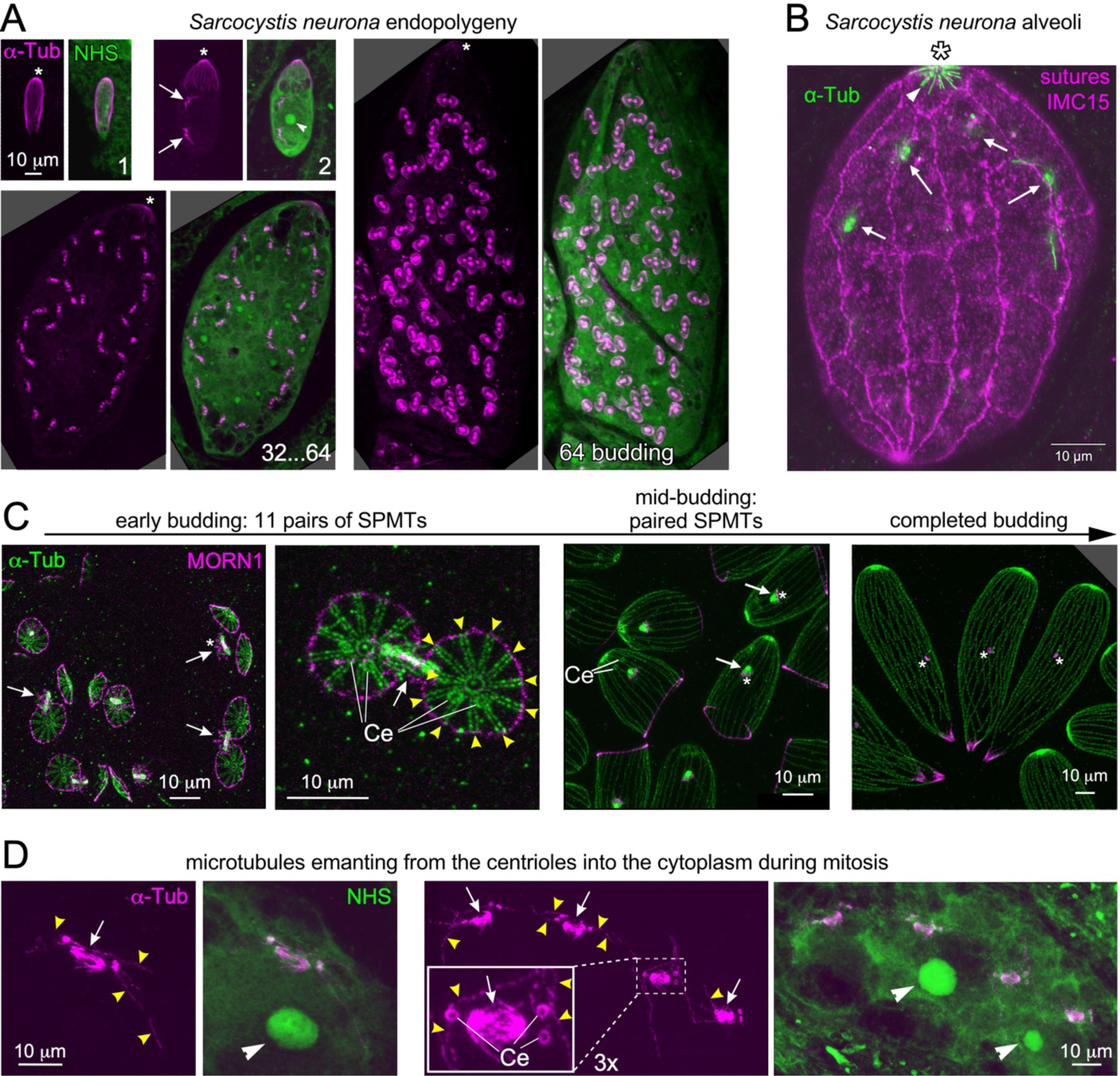
Spindle and SPMTs in *Sarcocystis neurona* endopolygeny. **A.** Representative stages of endopolygeny imaged by single expansion. 1: A just invaded, single merozoite; 2. Early schizont undergoing the first mitosis; 32-64: late schizont going through the last round of mitosis (32 spindles) coupled to budding 64 merozoites – the first outlines of the nSPMTs are visible at the ends of each spindle; 64 budding: later budding schizont with the nSPMTs well defined. Asterisks mark the apical end (conoid) of the mother’s cytoskeleton, which is visible till the very end; arrows mark mitotic spindles; arrowhead marks the nucleolus. All panels are from the same expansion experiment and are shown on the same scale. **B.** Single expansion imaging of the *S. neurona* IMC. The IMC is composed of an apical cap and 3×11 alveolar sheets (note that only the front of the schizont is displayed here). The sutures between the sheets are marked by stable overexpression of YFP-TgIMC15 [63]. Note the SPMTs (arrowhead) only extend along the apical cap in this mid-stage schizont. Arrows mark the mitotic spindles. **C.** Pan-ExM of *S. neurona* daughter budding stages co-stained with α-tubulin and MORN1. α-tubulin marks the spindle, centrioles and SPMTs. MORN1 highlights the centrocone and the basal complex. Ce: centriole; arrows mark mitotic spindle; yellow arrowheads mark the 11 pairs of nSPMTs; asterisks: centrocone. **D.** Pan-ExM of *S. neurona* schizonts, zoomed in on the spindles. MTs of unknown identity emanate from the centrioles into the cytoplasm during mitosis. Two different parasites are shown: left panels undergoing the first round of mitosis (two spindles); right panels undergoing third round of mitosis (eight spindles). Ce: centriole; arrows mark overlapping mitotic spindles; yellow arrowheads mark the MTs of unknown identity; white arrowheads mark the nucleolus.

Other distinctive observation on the *S. neurona* cortical cytoskeleton comprise the modest apical cap in schizonts, which is the only place where the mother’s SPMTs are seen (**Fig 3B**). In fact, during the progression of endopolygeny it appears that the mother’s SPMTs are shrinking (**Fig 3A**: all panels are on the same scale and from the same expansion experiment). This implies a putative level of catastrophe on the SPMTs (+)-ends, which goes against the current dogma of extreme SPMT stability [49]. Lastly, we very frequently observed MTs emanating from centrioles into the cytoplasm present at mitotic spindles in schizonts before the final round of S/M paired with daughter budding. Notably, bundled spindle MTs were present at the same time, but these were all clearly oriented toward the opposite centriole pair (**Fig 3D**). These cytoplasmically oriented MTs were somewhat reminiscent of astral MTs see in higher eukaryotes that function in orienting the mitotic spindle within the cell. Alternatively, this could be a leaky assembly of nSPMTs during mitotic rounds uncoupled from budding.

Overall, the comparative analysis of *S. neurona* as a parasite closely related to *T. gondii* highlighted several subtly distinctive features. Most notably that the paired nSPMT organization located at bud initiation corresponds directly with the number of alveolar membrane plates. Together this suggests an architectural layout rule, at least in the Coccidia, where nSPMTs raft organization directs alveolar plate organization.

### γ-Tubulin is critical for correct spindle and nSPMT assembly

Having established the early organization of nSPMTs, we next asked how the SPMTs are nucleated. To this end we examined the sub-cellular localization of γ-tubulin by pan-ExM. Using conventional microscopy, γ-tubulin was localized to the outer core of the centromeres, which is where the centrioles reside [18]. We endogenously tagged the C-terminus of γ-tubulin with the mini auxin inducible degron (mAID) coupled to five V5 epitopes (γ-tub-cKD). At the onset of mitosis, before budding starts, we observed γ-tubulin on and around the centrioles as well as the early mitotic spindle (**Fig 4A**). When the nSPMTs started forming, γ-tubulin resided on the spindle pole, the centrioles, and in a horseshoe shape between the forming conoid and the (−)-ends of the nSPMTs. γ-tubulin therefore seems to highlight the nAPR at bud initiation. During subsequent anaphase, when nSPMT rafts are fully formed, γ-tubulin is still present at the spindle pole and the centrioles, though with lower abundance, while it was absent from the apical end of the forming scaffold. We also detected γ-tubulin at the ICMTs within the conoid (**Fig 4A** panel 2). Thus, γ-tubulin is associated with the formation of all MT populations in tachyzoites. Furthermore, we sometimes see pairs of short γ-tubulin pairs emanating from a centriole (**Fig 4A**, panel 3 yellow arrowheads). We speculate that these could be the basis of new SPMTs pairs being initiated close to the centriole before they are transferred onto the nAPR.

**Fig. 4.**
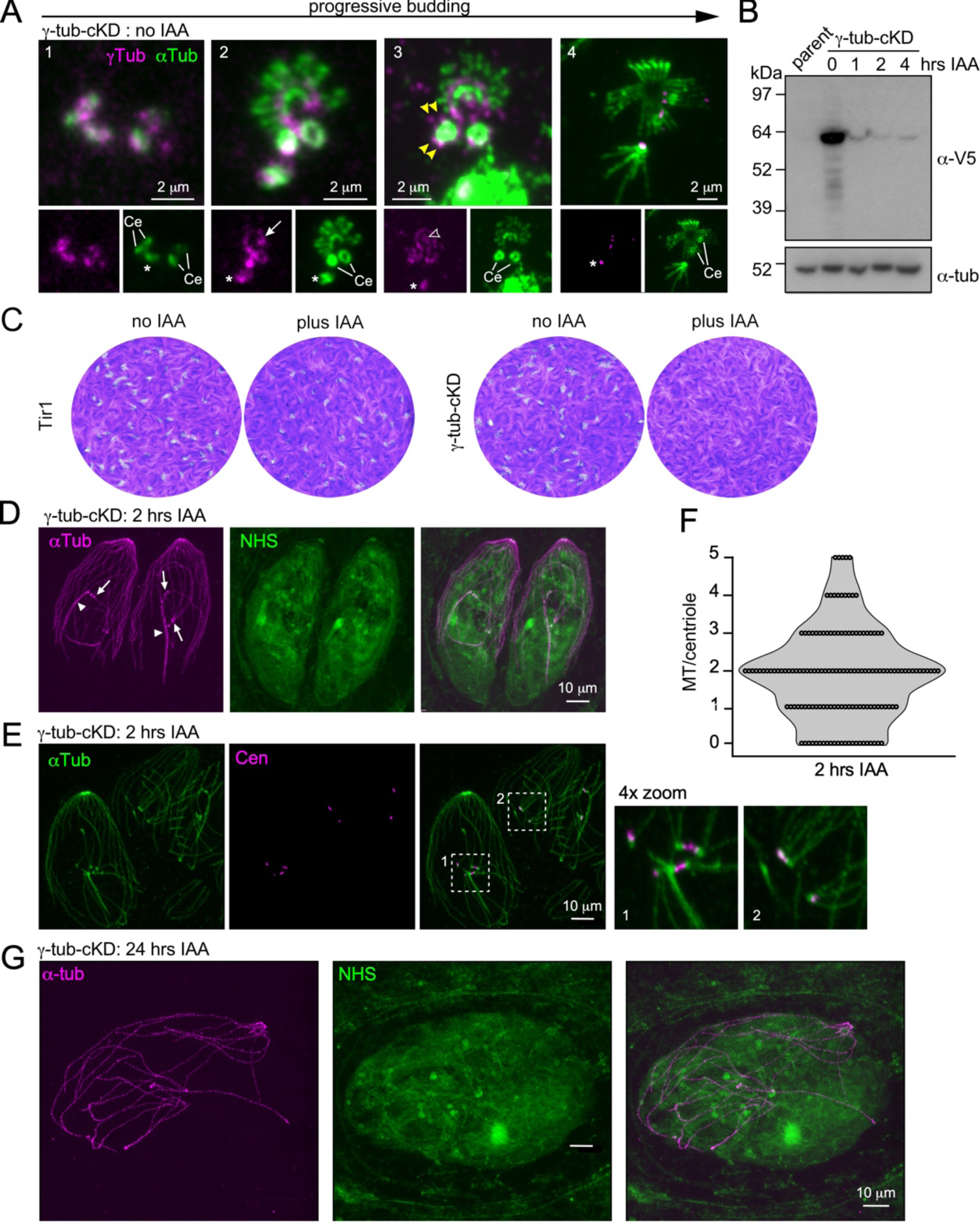
γ-tubulin is essential for cell division. **A.** Pan-ExM of parasites expressing γ-tubulin endogenously tagged at the C-terminus with mAID-5xV5 (γ-tub-cKD), co-stained with V5 and α-tubulin antisera. Asterisks: the spindle or spindle pole; arrow: horseshoe shape at the nAPR; open arrowhead: γ-tubulin at the ICMT; yellow arrowhead: γ-tubulin pairs of different length protruding from the centriole (Ce). **B.** Kinetics of γ-tub-cKD depletion upon IAA addition visualized by western blot. α-tubulin (MAb 12G10) is used as loading control. The mAID system allows fast protein degradation. **C.** Seven-day plaque assay of γ-tubulin depletion demonstrates its essentiality for the lytic cycle. **D.** Pan-ExM of 2 hrs γ-tubulin depleted parasites stained for α-tubulin and protein density marker NHS ester conjugated to ATTO488. Mother MT populations appear normal whereas two daughter MT populations are visible: one (arrows) connected to round structures (putatively centrioles), and the other (arrowheads) presents as a long bundle of MT. **E.** Pan-ExM of 2 hrs γ-tubulin depleted parasites co-stained for α-tubulin and *Hs*Centrin2 (centrosomes). A variable number of unstructured microtubules emanates from typically one side of each centriole, which are connected by Centrin. Numbered boxes are expanded in panels on the right. **F.** Quantification of panel D for the number of MTs emanating from centrioles. The majority of centrioles, roughly 30% is connected to 2 MTs, the remainder is a mixture of 0-5 MTs. 145 centrioles from 39 parasites were quantified. **G.** Depletion of γ-tubulin for 24 hrs imaged by pan-ExM reveals long unstructured MTs without any daughter cytoskeletons within an unstructured mother cell. NHS-ATTO488 reports on protein density.

To functionally dissect γ-tubulin, we took advantage of the mAID tag [64], facilitating effective depletion of most γ-tubulin within hours of indole-3-acetic acid (IAA) treatment (**Fig 4B**). When treated for seven days with IAA, γ-tubulin depleted parasites lost plaque forming capacity, demonstrating that it is essential to complete the lytic cycle (**Fig 4C**). Next, we assessed the effect of γ-tubulin depletion on the first round of daughter cell formation (2 hrs IAA), with a special focus on the MT cytoskeleton. The maternal SPMTs and conoid appeared unperturbed, but structures associated with daughter cell formation were severely disrupted. We observed two distinct MT populations; with one consisting of individual MTs connected to centrioles, the other formed by a bundle of MTs (**Fig 4D, E**). Centriole duplication and separation was not affected by γ-tubulin depletion, as indicated by co-staining with *Hs*Cen2 (**Fig 4E**), but the centrioles appeared denser, without their usually detected donut-like shape. We enumerated the variable number ‘centriolar’ MTs, which ranged from 0-5, with two MTs is the most common observed number (**Fig 4F**).

To test for consequences of γ-tubulin depletion beyond 2 hrs, we performed a 24 hr IAA treatment experiment, upon which the SPMTs of the mother are also perturbed (**Fig 4G**). During this period parasites failed to generate a new round of daughter scaffolds (**Fig 4G**, tubulin staining), however, the substantial size increase of the mother parasites suggested that many organelles still duplicated (**Fig 4G**, NHS).

To identify the observed MT populations and to assess fate of the spindle and nSPMTs in γ-tubulin depleted parasites, we stained with the IMC marker IMC6 [27, 65] and nSPMT marker polyglutamylated (PolyE) tubulin [22, 66]. This identified the ‘centriolar’ MTs as SPMTs, due to their association with the IMC and the presence of PolyE (**Fig 5A, B**). Notably, both markers did not stain the bundled MTs. Furthermore, MORN1 staining was seen in semi-circular structures surrounding the ‘centriolar’ MTs, likely indicating an incompletely formed BC. It further appeared to align with a subset of the ‘centriolar’ MTs (**Fig 5C**, γ-tub-cKD 2 hrs IAA). Collectively, the analysis of IMC6, PolyE, and MORN1 permitted a firm assignment of the ‘centriolar’ MTs as nSPMTs.

**Fig 5.**
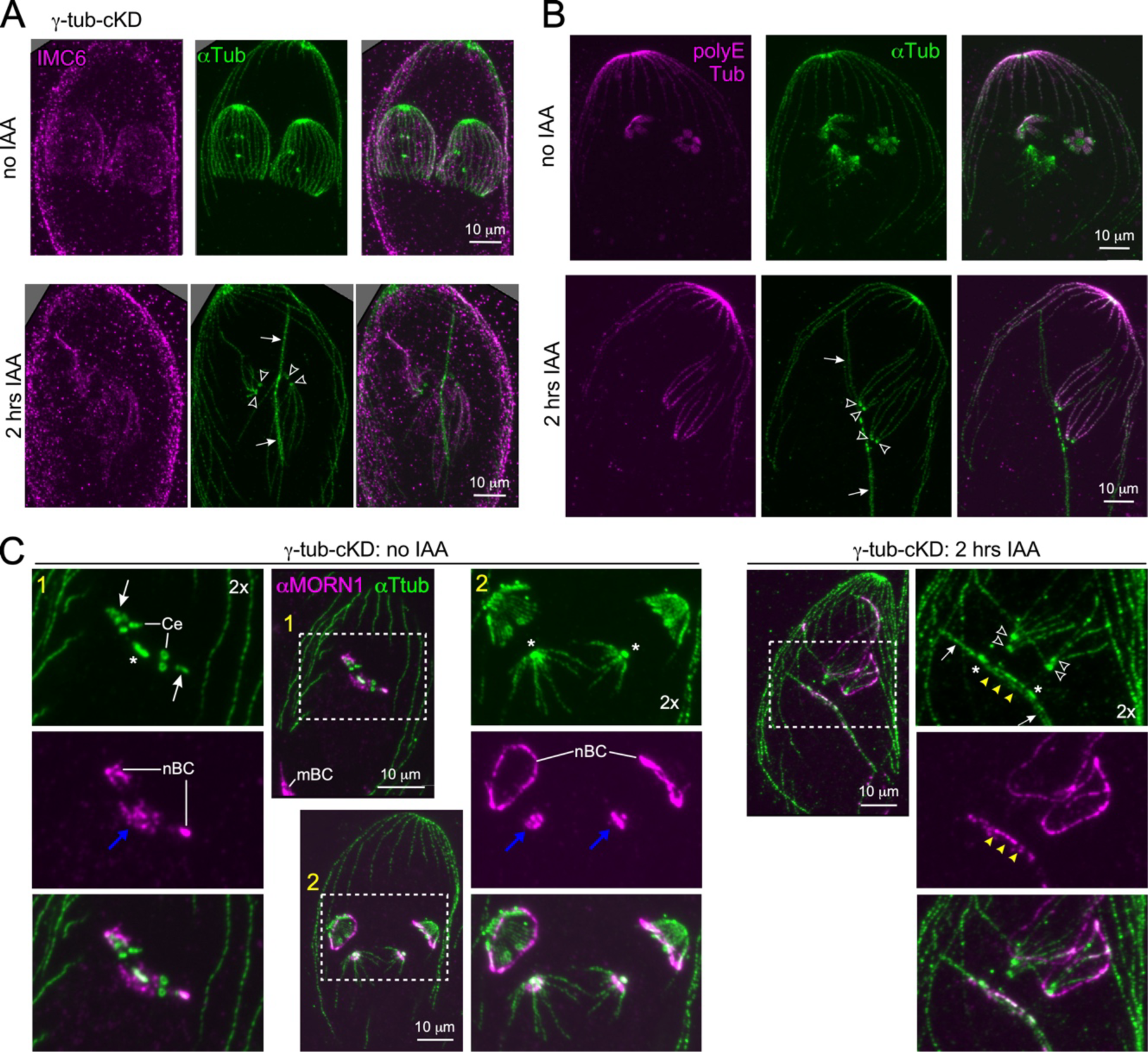
Identity of microtubule populations in γ-tubulin depleted parasites. **A.** Pan-ExM of control and 2 hrs γ-tubulin depleted parasites stained with α-tubulin and IMC6 antisera. Arrows: bundled MT, not coated with IMC6; open arrowheads: centrioles from which IMC6-coated MTs are emanating. **B.** Pan-ExM of control and 2 hrs γ-tubulin depleted parasites stained for α-tubulin and polyglutamylated (polyE) tubulin antisera. Arrows: bundled MT, not marked with polyglutamylated tubulin; open arrowheads: centrioles from which polyglutamylated tubulin-coated MTs are emanating. **C.** Pan-ExM of control and 2 hrs γ-tubulin depleted parasites stained for α-tubulin and MORN1 antisera. Yellow-1 labeled panels shows a very early budding cytoskeleton, with only a few nSPMTs (arrow) of the scaffold surrounded by the nascent basal complexes (nBC). MORN1 also highlights the centrocone (blue arrow) in the nuclear envelope, where the spindle reaches into the nucleus. Yellow-2 labeled panels show early budding parasites with dome shaped nSPMTs coated with the nBC close to the nSPMT (+)-ends. Blue arrows: centrocone. After γ-tubulin depletion (2 hrs IAA), MORN1 localizes in a semi-circle to the ‘centriolar” MTs and partially coats them. MORN1 also localizes to the middle of the bundled MT, highlighting the centrocone. mBC: mother parasite’s BC; asterisk: mitotic spindle; open arrowheads: centrioles; yellow arrowheads: bundled MTs coated with MORN1, marking the mitotic spindle embedded in the nuclear envelope. Dotted-line boxed areas are 2-fold enlarged. Yellow numbers indicate corresponding panels.

By extension of the above findings, the bundled MTs most likely represented the spindle MTs, which however, were much extended in length when compared to wildtype spindles. We confirmed this by use of MORN1, which besides the BC also marks the mitotic spindle at the centrocone, which is the structure embedded in the nuclear envelope through which the spindle MT are attached to the kinetochores (**Fig 5C**, panels 1 and 2) [33]. More saliently, MORN1 predominantly stained the bundled MTs in the middle (**Fig 5C**, yellow arrowheads), most likely highlighting the area where the spindle penetrates into the nucleus. The nature of MTs extending beyond the MORN1 stained bundle could either be extended spindle MTs or mis-oriented spindle MTs. Our current data cannot differentiate between these scenarios. In conclusion, γ-tubulin is critical for establishing all MT populations in tachyzoites, and germane to the MT daughter scaffold, facilitates correct nSPMT assemblage.

### γ-Tubulin depletion does not interfere with bud initiation

We recently promoted a model wherein the apical cap and the central and basal alveoli are assembled by different mechanisms from the BCC0 base, the former in the apical direction, and the latter in basal direction [21]. To further explore the state of the IMC in these parasites, and to determine the relationship between the nSPMT and assemblage of the IMC, we explored daughter assembly using the apical cap marker ISP1 [67] and the central and basal IMC marker IMC3 [27]. Interestingly, the apical cap’s appearance is unperturbed upon 2 hrs γ-tubulin depletion, forming apically of the duplicated centrosomes (**Fig 6A, B**). This indicated that this event in daughter bud formation does not require a correct assembly of the nSPMTs. In contrast, the assembly of the central and basal IMC highlighted by IMC3 is disorganized and results in nearly 100% misshapen daughter buds (**Fig 6C, D**). In conclusion, the data presented here supports our model wherein the apical cap is assembled by a mechanism different from the one governing the central and basal alveoli. Notably, correct nSPMT assembly is not required to form the apical but, is required for the normal formation of the central and basal alveolar IMC. These data consolidate the insights gleamed from the oryzalin experiment.

**Fig 6.**
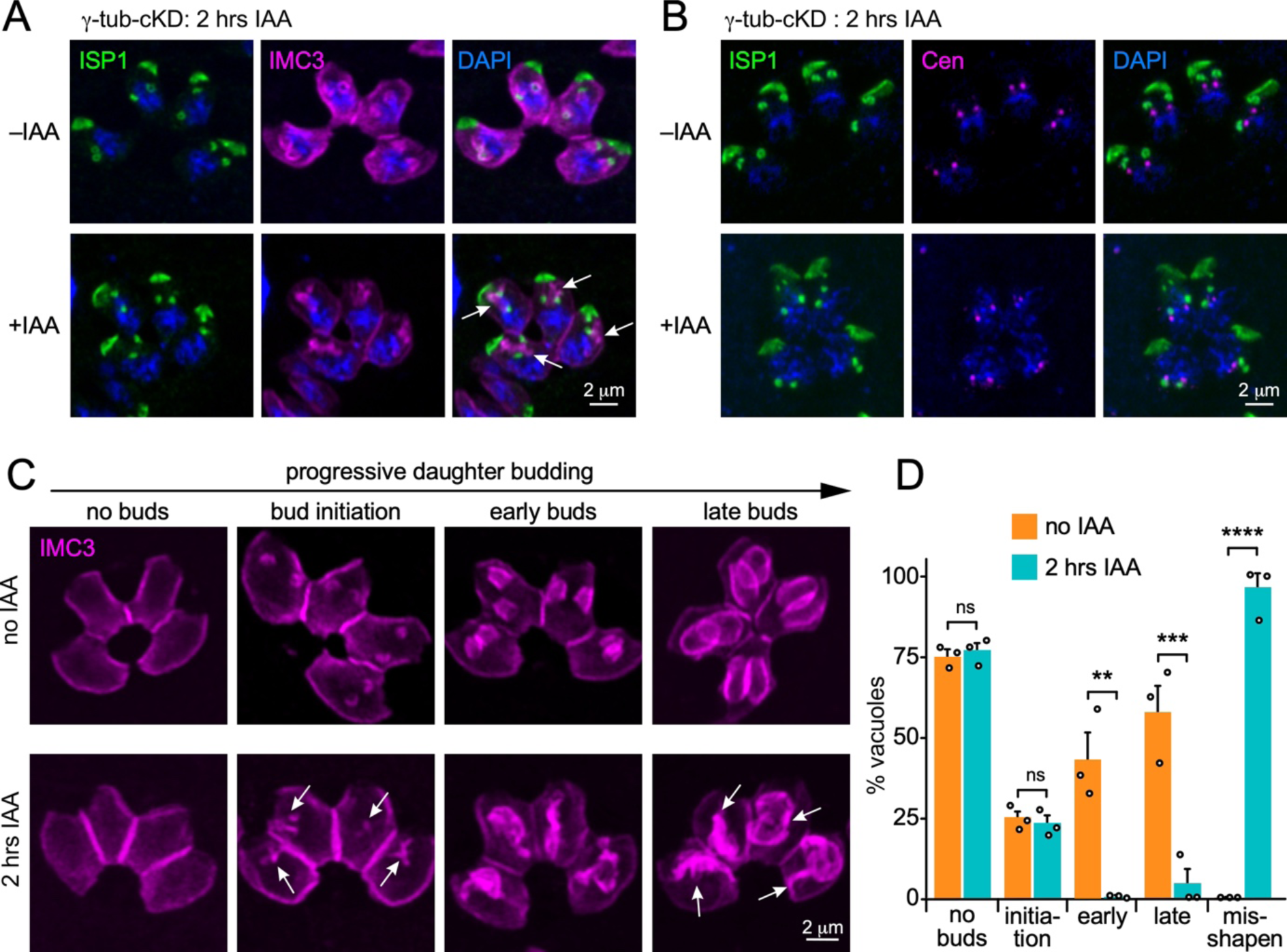
γ-tubulin depletion causes deviant daughter IMC buds. **A, B.** Airyscan superresolution images after 2 hrs of γ-tubulin depletion using two different co-stains: **A.** ISP1 (apical cap) and IMC3 (mother and daughter IMC); **B.** ISP1 (apical cap) and *Hs*Centrin2 (centrosomes). Arrows mark unstructured IMC3, whereas ISP1 appears normally clustered around the centrosomes. DAPI highlights DNA. **C.** Airyscan superresolution images of 2 hrs γ-tubulin depleted parasites. No interfereance with initiation of daughter budding is observed (arrows in ‘bud initiation’ panel) but the morphology of the daughter buds is severely affected: daughters appear disrupted and no cone-shape bud is assembled when γ-tubulin is depleted (arrows in ‘late buds’ panel). **D.** Quantification of panel C for incidence of budding and bud shapes upon 2 hrs of γ-tubulin depletion. n=3 biological replicates, 100 vacuoles each, error bars denote SEM. Data points indicate averages of individual replicates. One-way ANOVA with Tukey’s HSD. ns: not significant; ** *p* = 0.01; *** *p* = 0.001; **** *p* = 0.0001.

### SFA2 and γTuRC component GCP4 are different scaffolding factors

The SFA fiber arises from between the centriole pair and extends to the nascent apical complex of the daughter bud [19]. The current understanding of its function is positioning the MTOC, which forms the template for the daughter scaffold during cell division [19, 32]: SFA’s depletion disrupts daughter scaffold formation completely, assigning it an essential role the budding process [19]. We first re-examined the exact SFA2 localization here by the additional resolution offered by pan-ExM (**Fig 7A**). We confirmed the presence of SFA2 between the centriole pair very early in development, but in addition observe SFA2 in between the conoid and nSPTMs (panels 1 and 2). Once the bud transforms into the cone shape, during metaphase, SFA was seen as a fiber extending all the way from between the centrioles, to the apical side of the conoid. Notably, at the conoid, it appears as a flattened disk located above it and the nSPMTs (−)-ends (panel 3, arrowhead). Upon further budding, the SFA2 signal became discontinuous (panels 4) and upon bud maturation was only retained above the conoid (panel 5). Taken together, SFA2 is present near the nAPR during nSPMT formation, but progresses to a localization apical to the nascent scaffold.

**Figure 7.**
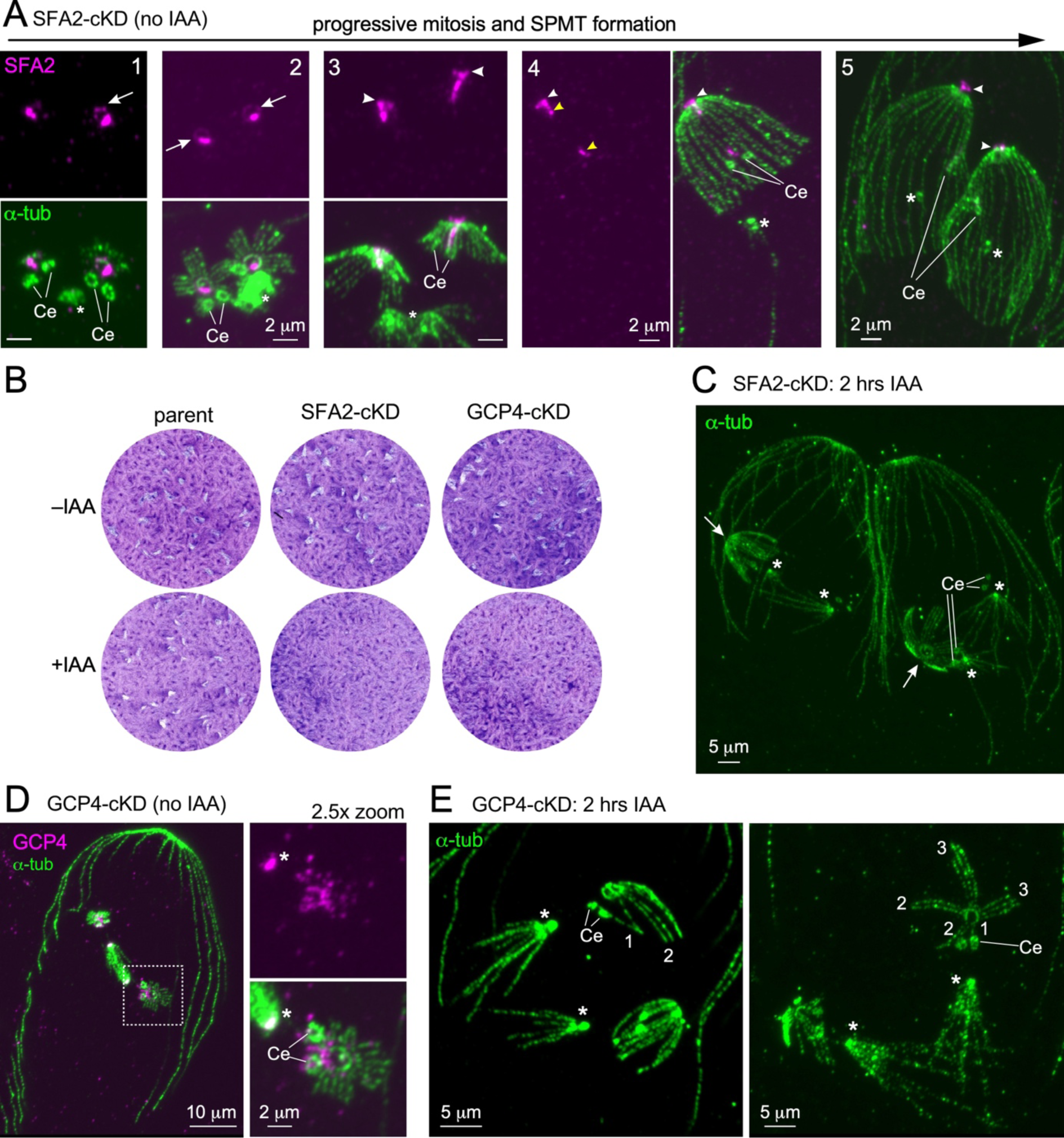
SFA2 and GCP4 depletion strongly affects bud formation. **A.** SFA2-mAID5xV5 (SFA2-cKD) localization throughout mitosis and early daughter bud assembly Ce: centriole; asterisks: mitotic spindle; arrows: SFA2 ring-like structure aligning with the conoid. Arrowhead: SFA2 signal apical to nAPR and conoid; yellow arrowhead mark the broken fiber at mid-budding. Imaged by pan-ExM. **B.** Seven-day plaque assays of SFA2 and GCP4 depleted parasites using the mAID system. Parent line is RHΔKu80-Tir1. **C.** Pan-ExM of 2 hr SFA2 depleted parasites. Mitosis progresses normally, but single daughter buds appear at a high frequency. The single buds display a normal nSPMT organization. Ce: centrioles; asterisks: mitotic spindle; arrowheads: apical end of the single daughter bud. **D.** GCP4-mAID-5xV5 (GCP4-cKD) localizes to the spindle poles (asterisks) as well as in between the centrioles, and to the forming scaffold. Imaged by pan-ExM. **E.** Pan-ExM of 2 hr GCP4 depleted parasites. Mitosis progresses normally, and two daughter buds are formed, however, the organization and number of nSPMTs in each sheet is irregular (number of nSPMTs per sheet indicated by numericals). Ce: centrioles; asterisks: spindle poles.

Next, we depleted SFA2 using the mAID system, which resulted in non-viable parasites (**Fig 7B**). A short, 2 hrs SFA2 depletion resulted in normal appearing centriole pairs and mitotic spindles, but parasites formed only a single daughter nSPMTs scaffold, which was seen at a high frequency (**Fig 7C**). The nSPMT constellation remarkably was the same as seen in unperturbed parasite buds. The mechanism seems to be an all or nothing, i.e. SFA2 depletion below a certain threshold results in no daughter MT scaffold, whereas above this threshold a normal scaffold is assembled. Notably, we did not detect any ‘centriolar’ SPMTs, as seen after γ-tubulin depletion (**Fig 4&5**), in absence of SFA2.

Together, we confirm SFA2’s previously reported essential role of establishing the daughter scaffold [19].

The essential involvement of γ-tubulin in scaffold formation however, led us to further pursue the function of the γ-tubulin ring complex (γTuRC). *T. gondii* encodes all γ-TuRC members ([52]; with GCP5 (TGME49_288890) and GCP6 (TGME49_285780) recently being annotated on the ToxoDB genome database [68]). Here, we explored the role of γ-TuRC using a conditional GCP4-mAID-5xV5 (GCP4-cKD) parasite line. Firstly, it was observed that GCP4 is essential for the lytic cycle as no plaques form upon its depletion (**Fig 7B**). When we analyzed its subcellular localization via pan-ExM, GCP4 was present at the spindle poles, in between the centrioles, and at the early scaffold (**Fig 7D**). These are all localizations that were also observed for γ-tubulin (**Fig 4A**). After 2 hrs of GCP4 depletion, daughter buds were strongly affected and exhibited disorganized nSPMTs (**Fig 7E**). Here, nSPMT rafts contained less than the usually four nSPMTs, but were formed by either one, two, or three nSPMT per raft (**Fig 7E**, numericals). Interestingly, the spindle and the centrioles appeared unaffected by GCP4 depletion under the given condition, although GCP4 localizes to both structures.

In summary, our data highlight the essential involvement of γ-tubulin and γ-TuRC in daughter scaffold formation.

## Discussion

Here, we will put our new findings in chronological order to reconstruct the series of sequential events underlying cell division and how the formation of the different structural components is hierarchically organized as summarized in **Fig 8**.

**Figure 8.**
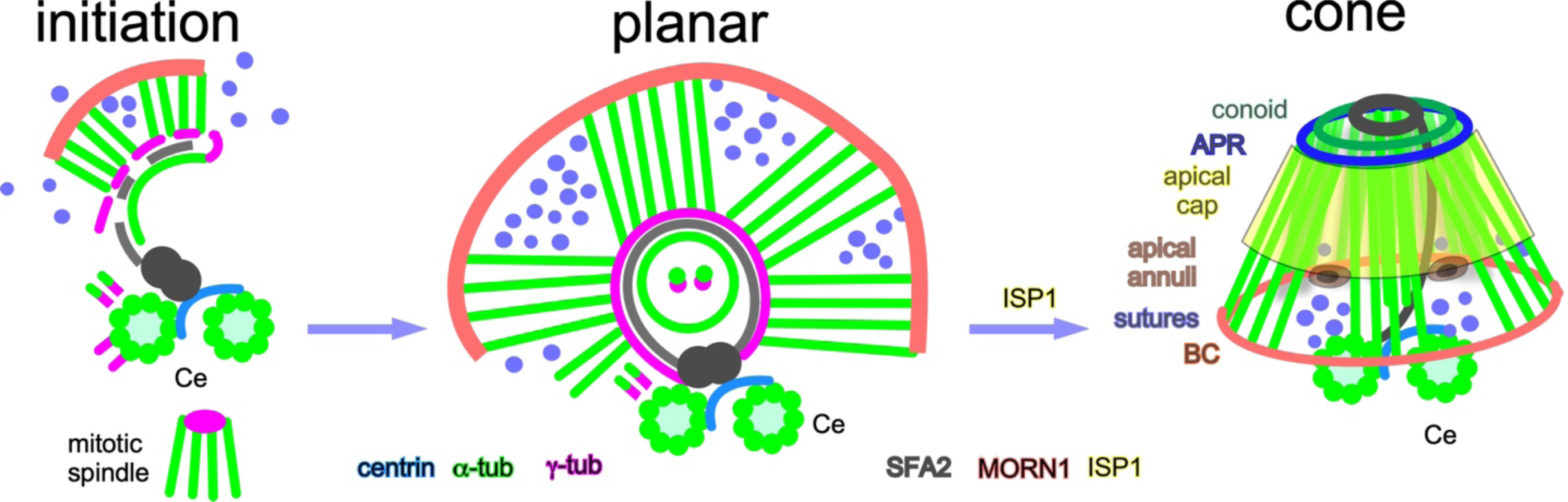
Model of daughter scaffold formation in *T. gondii*. Summarizing schematic: The daughter scaffold is initiated close to the centrioles (Ce) and relies on two factors; γ-tubulin (magenta) for the nucleation and/or stabilization/anchoring of tubulin components, and as previously reported on the SFA fiber [19] (black). The basal complex (MORN1 [31], red), the alveoli (not shown), and IMC components (e.g. BCC0 [21] and IMC32 [23], blue) align with the earliest observable tubulin scaffold structure. SPMTs are nucleated close to the centrioles, presumably in the pericentriolar material of the outer centrosomal core, and subsequently added to the forming APR (highlighted by γ-tubulin/magenta). Closure of the circular conoid and nAPR, in concert with nSPMT addition, establishes the scaffold’s planar configuration. At this point the basal complex assumes ring shape. Most abundantly, four rafts of four nSPMTs and one raft of six nSPMTs (this study and [56]) support the alveoli, which are being added to the growing scaffold (not shown), The apical cap (ISP1 [67], yellow) is formed by a so far incompletely understood process, but presumably establishes the cone shape of the scaffold, which then growths in basal direction. At this point γ-tubulin no longer localizes to the APR (blue), which is then stabilized by other factors (e.g. KinA/APR1 [42] and/or AC9/10 [22]).

First of we would like to point out a potential caveat in our experiment design: the very short depletion window of 2 hrs we used, which is needed to capture the earliest events in the developing phenotypes. Since efficient methods to synchronize *T. gondii* cell division to the degree needed for these experiments have not been established, we cannot be sure that replication did start in the complete absence of any of the proteins we targeted. In addition, the mAID system does not necessarily completely remove all protein, but it’s extremely fast kinetics of depletion makes it the best tool for this study. Either way, putative protein traces can influence the phenotypes we observed.

We define a key role for γ-tubulin, as it is present at the formation site of all the tachyzoite’s MT populations: spindle in the centrosome inner-core, centrioles in the centrosome outer-core, the nSPMTs surrounding the conoid, and the ICMTs (**Fig 4**). Pertinent to daughter scaffold assembly, there are key observations regarding the positioning of γ-tubulin in the extending horseshoe shape emanating from the centriole, which we interpret as the nAPR. As the first sign of the scaffold formation, we observe a short, single MT coated with MORN1 (**Fig 5C**, arrows in panel 1). The assembly then progresses by forming nSPMTs in close proximity to the centrioles, as we see paired γ-tubulin (**Fig 4A**) as well as α-tubulin appendages sprouting from there (**Fig 2B**). These novel paired MT assemblies then apparently transfer from the centrosome to the growing APR, which contains both γ- and α-tubulin, although the mechanics of this step cannot be fully deduced from the images we captured. Supportive data for MT nucleation close to the centrioles are found in *S. neurona* where we observe MTs associated with centrioles as well (**Fig 3D**). Although their function remains unknown for now, their presence suggests that the outer core of the centrosome can nucleate MTs.

In the above model, nSPMT initiation close to the centrioles precedes their transfer to the nAPR. Upon γ-tubulin depletion we find the daughter scaffold entirely absent, indicating that γ-tubulin is essential for its formation. The absence of the scaffold likely explains the observed ‘centriolar’ SPMTs phenotype (**Fig 5**), visible after γ-tubulin depletion, as SPMTs fail to transfer to the nAPR and remain associated with their nucleation place. Based on the timeframe of experiments and the depletion system used, we can however, not exclude that SPMT nucleation also depends on γ-tubulin.

We further dissected if and how γ-TuRC component GCP4 acts in this process by examining its localization pattern and phenotype upon its depletion. This showed that GCP4 is also at the nAPR and although its depletion does not mirror the γ-tubulin phenotype, it severely affects the efficiency of nSPMT rafts assembly. What is curious though is that even if one nSPMT assembly is skipped, the SPMT grouping in rafts proceeds as if the nSPMT was assembled. As such, γ-TuRC is critical for correct nSPMT raft architecture and may facilitate specificity of γ-tubulin localization and or function at the forming scaffold. We currently do not know how γ-tubulin is differentially targeted to a variety of structures but note that all of them are located closely together and are formed at the same time, suggesting differences in the composition of the respective γ-TuRCs. To gain the necessary fine-tuning, γ-TuRCs could for example rely on post-translational modifications, and GCP2 was recently identified as a substrate of TgCrk4, a kinase that acts upstream of the spindle assembly and centrosome reduplication checkpoint [69].

Here, we also explored the connection between the number of nSPMTs rafts and the architecture of the alveolar plate organization in the IMC. TEM demonstrated the presence of membrane on top the SPMT rafts (**Fig 2C**), which is in line with a few other published TEM micrographs revealing the organization of the IMC around five-fold nSPMT raft symmetry [70, 71] and a more recent Cryo-ET study [57]. The key observation is the discontinuity between the neighboring membrane sheets, which we infer is where the longitudinal sutures between the alveolar plates are positioned. We employed BCC0 and IMC32 as markers of this alveolar architecture. Depolymerizing nSPMT via oryzalin demonstrates that the symmetrical deposition of the longitudinal sutures is lost, which puts nSPMT raft organization before putting down the IMC sutures. Furthermore, the 11 alveolar plates in *S. neurona* corresponded directly with nSPMT rafts composed of two nSPMTs (**Fig 3**). Thus, across two different Coccidian parasites the deposition of nSPMT rafts cues the number of alveolar plates. Furthermore, there is a strong proclivity to form paired MTs observed in both *T. gondii* and *S. neurona*, as well as upon γ-tubulin depletion (**Fig 2A, 4F**). Whether these rules apply universally across all Apicomplexa is unlikely, since different ‘zoites’ display vastly different alveolar quilts and SPMT numbers [2], but it at least applies to the Coccidia.

A distinct feature in the Coccidia is the presence of a single apical cap alveolus, separated from the central and basal alveolar plates. Upon depletion of γ-tubulin we learned that apical cap formation (as marked by membrane anchored ISP1 [67]) is independent from nSPMT formation, but that the correct assembly of the central and basal rows of alveoli does require the nSPMTs (**Fig 5**). This aligns with previous observations using oryzalin, demonstrating the nSPMTs are not required for ISP1 deposition [67]. Moreover, we have recently shown that the basis for the IMC is the five-fold BCC0 organization, from which the apical cap is assembled in the apical direction, and the central and basal alveolar plates in the basal direction (**Fig 1A**).

At the mid-budding stage, the nSPMT raft organization transitions into an equal distribution of nSPMTs around the APR. Comparison of *T. gondii* tachyzoites with *S. neurona* merozoites indicates this could be tailored to individual needs, since even in completely formed *S. neurona* merozoites the SPMTs still have a paired appearance (**Fig 3C**). A recent study in *T. gondii* found that equal distribution of nSPMT coincides with the alignment of APR1 to the APR [56]. Although mechanistically, we currently have no insights into how the distribution, or maintenance of clustering, is organized. Another distinctive, and putative related, feature is the differential length of the SPMTs: in *T. gondii* they are largely of similar length running to about 2/3 the length of the tachyzoite, whereas in *S. neurona* merozoites the pairs consist of one long MT running all the way to the basal complex, and one MT ending at about 2/3 the length of the merozoite (**Fig 3C**). Clearly, there is very precise control of the SPMT length, which could be related to their raft organization.

Overall, the pairing of pan-ExM with genetic dissection of early SPMT formation and its interface with other cytoskeletal elements has delivered several advancing new insights regarding the interdependent assembly of the different apicomplexan cytoskeleton elements. We also uncovered a new correlation between SPMT and alveolar architecture. However, many questions remain, e.g. how the parasite counts SPMTs, and how they transition from the centrosome to the nAPR. The insights and tools developed here raised these new questions and at the same provided the tools needed toward answering them.

## Material and Methods

### Parasites and host cell cultures

All *T. gondii* strains are derived from RH, and tachyzoite cultures were maintained in human telomerase reverse transcriptase (hTERT)-immortalized human foreskin fibroblasts (HFFs) as previously described [72]. Immunofluorescence assays (IFAs) were performed using primary HFFs. The *Sarcocystis neurona* SN3ΔHXGPRT-G11 strain (kindly shared by Dr. Daniel Howe, University of Kentucky) was maintained and imaged in bovine turbinate (BT – ATCC CRL-1390^™^) cell monolayers described in [73, 74], supplemented with a chemically defined lipid concentrate at a 1:1000 dilution according to manufacturer protocol (Gibco™ 11905031). *S. neurona* stably transfected with tub-TgIMC15-eYFP to visualize parasite IMC sutures [63] were generated under 10 μM pyrimethamine selection and cloned by limiting dilution [73]. Conditional depletion lines were established by targeting the endogenous 3’end of γ-tubulin (TGGT1_226870A+B), SFA2 (TGGT1_205670) and GCP4 (TGGT1_209970) with a mAID-5xV5-DHFR homologous repair construct, after CRISPR/Cas9 induced DNA double strand break in the respective region. Selection for stable *T. gondii* transfectants was performed under 1 µM pyrimethamine and stable lines were cloned by limiting dilution (γ-tubulin-cKD and GCP4-cKD). Protein knock-downs using the mAID system were performed under 500 µM auxin (IAA: 3-indoleacetic acid in 100% ethanol; Sigma-Aldrich) [64].

### (Immuno-) fluorescence microscopy

Intracellular parasites grown overnight in a 6-well plate containing coverslips confluent with HFF cells were fixed with 100% methanol and blocked for 30 min with 2 % BSA in 1x PBS. The primary antisera (IMC3 and *Hs*Centrin2 [21], 1:1,000; ISP1 [67[ 1:500) and secondary antisera (Thermo Fisher Scientific; goat anti-mouseALEXA488 and goat anti-rabbitALEXA594, 1:500) were diluted in blocking solution and applied for 30 min at RT, followed by three 5 min washes in 1x PBS. DNA staining was performed using 4’,6-diamidino-2-phenylindole (DAPI) in the first wash after the last antibody.

### Iterative expansion microscopy (pan-ExM)

Pan-ExM of *T. gondii* tachyzoites and *S. neurona* schizonts/merozoites was achieved by following published protocols [53, 54]. Briefly, tachyzoites were grown over night in 6-well dishes on confluent HFF cells, treated for indicated time frames with auxin (IAA) and fixed with 4% PFA. *S. neurona* merozoites were seeded on confluent BT cells and grown up to120 hrs before fixation with 4% PFA in PBS. Samples were incubated in anchoring solution (0.7% Formaldehyde, 1% Acrylamide in PBS) for 3-4 hrs at 37°C. To achieve a consistent primary gel thickness, samples were placed in gelation chambers [53, 75] and incubated with primary gel solution (19% sodiumacrylate, 10% acrylamide, 0.1% DHEBA in PBS, 0.25% TEMED and APS) for 1 hr at 37°C. Resulting gels were first denaturized in a buffer (200 mM SDS, 200 mM NaCl, 50 mM Tris, pH 6.8) for 15 min at RT and subsequently incubated at 73°C for 1 hr. Gels were washed in PBS and left for first expansion overnight in ultrapure water. At the next day, a 2×2 cm gel piece was taken for further processing and incubated in neutral hydrogel solution (10% acrylamide, 0.05% DHEBA, in H_2_O, 0.05% TEMED and APS), twice –each time for 20 min– at RT. Excess hydrogel solution was carefully removed with a Kim Wipe before gels were sandwiched between a microscopy slide and a 22 mm coverslip, followed by an 1 hr incubation at 37°C. Gels were then briefly washed in PBS and incubated twice –each time for 15 min on ice– in secondary gelation solution (19% sodiumacrylate, 10% acrylamide, 0.1% bis-acrylamide in PBS, 0.05% TEMED and APS). Excess gelation solution was carefully removed with a Kim Wipe, gels sandwiched between a microscopy slide and a 22 mm coverslip, and incubated for 1 hr at 37°C. To dissolve DHEBA, gels were placed in 0.2 M NaOH in H_2_O for 1 hr at RT and subsequently washed in PBS until the pH was neutral.

Pieces of the final gel were used for antibody staining. Primary antibodies (rabbit anti-V5 (Abcam) 1:300; rabbit anti-MORN1 [31] 1:500; rabbit anti-PolyE (AdipoGen) 1:500; rabbit anti-IMC6 [65] 1:500; mouse anti-α tubulin (clone 12G10, DSHB) 1:250, and rabbit anti-GFP (Torrey Pines Biolabs) 1:300) were diluted in 2% BSA in PBS and gels incubated overnight at RT, followed by a 3 hrs incubation step at 37°C at the next day. Gels were then extensively washed in 0.1% PBST before incubated in secondary antibody solution (Thermo Fisher Scientific; goat anti-mouseDyLight488, goat anti-rabbitDyLight594, goat anti-mouseALEXA488, goat anti-rabbitALEXA594, all 1:200). Incubation times and temperatures mirrored conditions used for primary antibodies and gels were washed in 0.1% PBST. To stain for protein density, gels were incubated in NHS-ester-ATTO488 (20 µg/mL in PBS) for one hour and subsequently washed with 0.1% PBST. The final expansion was done in in ultrapure water overnight. For imaging, expanded gels were placed in 35 mm glass bottom dishes (Ibidi) coated with poly-L Lysine and imaged on a ZeissLSM880 microscope with Airyscan unit. Images were acquired with ZeissZen Black software, while ZeissZenBlue version 2.3 was used for image processing using standard settings. Final image processing was done in FIJI [76] by adjusting brightness and contrast.

### Single expansion of *Sarcocystis neurona* samples

Single expansion of *Sarcocystis* samples was done similar to a previously reported protocol [56]. Briefly, the pan-ExM protocol was followed until the first expansion in ultra-pure water over night. Pieces of the gel were then incubated in PBS to allow gel shrinkage and subsequently stained with primary antibodies in 2% BSA/PBS overnight. Gels were washed in 0.1% PBST and subjected to secondary antibody staining overnight. Antibody dilutions were the same as used for pan-ExM. At the next day, gels were extensively washed with 0.1 % PBST. To stain for protein density, gels were incubated in NHS-ester-ATTO488 (5 µg/mL in PBS) for one hour and subsequent washed with 0.1% PBST. The final expansion was done in ultrapure water overnight and gels were imaged the next day as stated in the pan-ExM section.

### Transmission electron microscopy (TEM)

Wild type RH parasites were grown overnight in a T25 tissue culture flask confluent with HFF cells. The intracellular parasites were collected by trypsinizing the monolayer and fixed with 2.5% glutaraldehyde in 0.1 M phosphate buffer pH 7.2. The samples were fixed in OsO_4_, dehydrated in ethanol, treated with propylene oxide and embedded in Spurr’s epoxy resin. Sections were stained with uranyl acetate and lead citrate prior to examination in a JEOL 1200 EX electron microscope [77].

### Chemical perturbation of MT with oryzalin

Oryzalin treatment was done similar to previous reports [60] [61] *.T. gondii* tachyzoites expressing endogenously tagged IMC32-5xV5 were grown in 6-well plates, containing coverslips with confluent HFF cells, overnight. At the next day, parasites were treated with a final concentration of 2.5 µM oryzalin in DMSO and incubated for 2 hrs under standard culture conditions. Untreated IMC32-5xV5 expressing parasites were used as controls. Parasites were fixed with 4% PFA in PBS and subjected to the pan-ExM protocol.

### Western blotting

For western blotting, lysates of 2×10^6^ parasites (either IAA treated/not treated γ-Tub-cKD parasites or RHΔKu80-Tir1) were loaded on/separated via SDS-PAGE and proteins transferred on PVDF membranes. Membranes were blocked in 6% milk in PBS and subsequently stained with mouse anti-V5 (clone SV5-Pk1, BioRad, 1:10 000) and goat anti-mouse (1:100 000) antibodies. The secondary antibody is coupled to HRP and the Super Signal West Atto Ultimate Sensitivity Substrate (Thermo Scientific) was used as a HRP substrate. The membrane was stripped of antibodies and mouse anti α-tubulin (12G10, DSHB, 1:15 000) antibody was used as loading control.

## Acknowledgements

We would like to thank VEuPathDB for the readily accessible *T. gondii* genome assemblies and their functional annotations, Dr. Daniel Howe, University of Kentucky, for sharing reagents, Dr. Peter Bradley, University of California, Los Angeles, for ISP1 and IMC6 antisera, Dr. Iain Cheeseman, Massachusetts Institute of Technology, for *Hs*Cen2 antiserum, Garret Eichlin and John Cook for tireless cloning efforts and Dr. Jennifer Wang, Washington University, St. Louis for helpful discussions. Special thanks to Bret Judson and the Boston College Imaging Core for infrastructure, support, and discussions.

This study was supported by the National Institute of Health (NIH) grants R21AI171821 and R21AI180525 to KE, American Heart Association (AHA) award 1014105 to CB, R01AI152387 to MJG, and a Major Research Instrumentation grant 1626072 from the National Science Foundation (NSF). The funders had no role in study design, data collection and analysis, decision to publish, or preparation of the manuscript.

## References cited

1. Gubbels, M.J., C.D. Keroack, S. Dangoudoubiyam, H.L. Worliczek, A.S. Paul, C. Bauwens, B. Elsworth, K. Engelberg, D.K. Howe, I. Coppens, and M.T. Duraisingh, 2020. Fussing About Fission: Defining Variety Among Mainstream and Exotic Apicomplexan Cell Division Modes. Front Cell Infect Microbiol, 10: 269.

2. Harding, C.R. and F. Frischknecht, 2020. The Riveting Cellular Structures of Apicomplexan Parasites. Trends Parasitol, 36: 979–991.

3. Harding, C.R. and M. Meissner, 2014. The inner membrane complex through development of Toxoplasma gondii and Plasmodium. Cell Microbiol, 16: 632–41.

4. Gubbels, M.J. and N.S. Morrissette, The Toxoplasma cytoskeleton: structures, proteins, and processes, in Toxoplasma gondii: the model apicomplexan - perspective and methods, L.M. Weiss and K. Kim, Editors. 2020, Academic Press: London, IK. 743–788.

5. Goodenough, U., R. Roth, T. Kariyawasam, A. He, and J.H. Lee, 2018. Epiplasts: Membrane Skeletons and Epiplastin Proteins in Euglenids, Glaucophytes, Cryptophytes, Ciliates, Dinoflagellates, and Apicomplexans. MBio, 9: e02020–18.

6. Anderson-White, B.R., J.R. Beck, C.T. Chen, M. Meissner, P.J. Bradley, and M.J. Gubbels, 2012. Cytoskeleton assembly in Toxoplasma gondii cell division. Int. Rev. Cell Mol. Biol., 298: 1–31.

7. Mann, T. and C. Beckers, 2001. Characterization of the subpellicular network, a filamentous membrane skeletal component in the parasite Toxoplasma gondii. Mol Biochem Parasitol, 115: 257–68.

8. Gould, S.B., W.H. Tham, A.F. Cowman, G.I. McFadden, and R.F. Waller, 2008. Alveolins, a new family of cortical proteins that define the protist infrakingdom Alveolata. Mol Biol Evol, 25: 1219–30.

9. Al-Khattaf, F.S., A.Z. Tremp, and J.T. Dessens, 2015. Plasmodium alveolins possess distinct but structurally and functionally related multi-repeat domains. Parasitol Res, 114: 631–9.

10. Russell, D.G. and R.G. Burns, 1984. The polar ring of coccidian sporozoites: a unique microtubule-organizing centre. J Cell Sci, 65: 193–207.

11. Sun, S.Y., L.A. Segev-Zarko, M. Chen, G.D. Pintilie, M.F. Schmid, S.J. Ludtke, J.C. Boothroyd, and W. Chiu, 2022. Cryo-ET of Toxoplasma parasites gives subnanometer insight into tubulin-based structures. Proc Natl Acad Sci U S A, 119.

12. Dos Santos Pacheco, N., N. Tosetti, L. Koreny, R.F. Waller, and D. Soldati-Favre, 2020. Evolution, Composition, Assembly, and Function of the Conoid in Apicomplexa. Trends Parasitol, 36: 688–704.

13. Tomasina, R., F.C. Gonzalez, S. Echeverria, A. Cabrera, and C. Robello, 2023. Insights into the Cell Division of Neospora caninum. Microorganisms, 12.

14. D’Haese, J., H. Mehlhorn, and W. Peters, 1977. Comparative electron microscope study of pellicular structures in coccidia (Sarcocystis, Besnoitia and Eimeria). Int J Parasitol, 7: 505–18.

15. Porchet, E. and G. Torpier, 1977. Freeze fracture study of Toxoplasma and Sarcocystis infective stages. Z Parasitenkd, 54: 101–24.

16. Francia, M.E. and B. Striepen, 2014. Cell division in apicomplexan parasites. Nature reviews. Microbiology, 12: 125–36.

17. Chen, C.T. and M.J. Gubbels, 2013. The Toxoplasma gondii centrosome is the platform for internal daughter budding as revealed by a Nek1 kinase mutant. J Cell Sci, 126: 3344–55.

18. Suvorova, E.S., M. Francia, B. Striepen, and M.W. White, 2015. A novel bipartite centrosome coordinates the apicomplexan cell cycle. PLoS Biol, 13: e1002093.

19. Francia, M.E., C.N. Jordan, J.D. Patel, L. Sheiner, J.L. Demerly, J.D. Fellows, J.C. de Leon, N.S. Morrissette, J.F. Dubremetz, and B. Striepen, 2012. Cell division in Apicomplexan parasites is organized by a homolog of the striated rootlet fiber of algal flagella. PLoS biology, 10: e1001444.

20. Tomasina, R., F.C. Gonzalez, and M.E. Francia, 2021. Structural and Functional Insights into the Microtubule Organizing Centers of Toxoplasma gondii and Plasmodium spp. Microorganisms, 9.

21. Engelberg, K., T. Bechtel, C. Michaud, E. Weerapana, and M.J. Gubbels, 2022. Proteomic characterization of the Toxoplasma gondii cytokinesis machinery portrays an expanded hierarchy of its assembly and function. Nat Commun, 13: 4644.

22. Tosetti, N., N. Dos Santos Pacheco, E. Bertiaux, B. Maco, L. Bournonville, V. Hamel, P. Guichard, and D. Soldati-Favre, 2020. Essential function of the alveolin network in the subpellicular microtubules and conoid assembly in Toxoplasma gondii. Elife, 9: e56635.

23. Torres, J.A., R.R. Pasquarelli, P.S. Back, A.S. Moon, and P.J. Bradley, 2021. Identification and Molecular Dissection of IMC32, a Conserved Toxoplasma Inner Membrane Complex Protein That Is Essential for Parasite Replication. mBio, 12.

24. Baptista, C.G., A. Lis, B. Deng, E. Gas-Pascual, A. Dittmar, W. Sigurdson, C.M. West, and I.J. Blader, 2019. Toxoplasma F-box protein 1 is required for daughter cell scaffold function during parasite replication. PLoS Pathog, 15: e1007946.

25. O’Shaughnessy, W.J., X. Hu, T. Beraki, M. McDougal, and M.L. Reese, 2020. Loss of a conserved MAPK causes catastrophic failure in assembly of a specialized cilium-like structure in Toxoplasma gondii. Mol Biol Cell, 31: 881–888.

26. Back, P.S., A.S. Moon, R.R. Pasquarelli, H.N. Bell, J.A. Torres, A.L. Chen, J. Sha, A.A. Vashisht, J.A. Wohlschlegel, and P.J. Bradley, 2023. IMC29 Plays an Important Role in Toxoplasma Endodyogeny and Reveals New Components of the Daughter-Enriched IMC Proteome. mBio, 14: e0304222.

27. Anderson-White, B.R., F.D. Ivey, K. Cheng, T. Szatanek, A. Lorestani, C.J. Beckers, D.J. Ferguson, N. Sahoo, and M.J. Gubbels, 2011. A family of intermediate filament-like proteins is sequentially assembled into the cytoskeleton of Toxoplasma gondii. Cell Microbiol, 13: 18–31.

28. Harding, C.R., M. Gow, J.H. Kang, E. Shortt, S.R. Manalis, M. Meissner, and S. Lourido, 2019. Alveolar proteins stabilize cortical microtubules in Toxoplasma gondii. Nat Commun, 10: 401.

29. Back, P.S., W.J. O’Shaughnessy, A.S. Moon, P.S. Dewangan, M.L. Reese, and P.J. Bradley, 2022. Multivalent Interactions Drive the Toxoplasma AC9:AC10:ERK7 Complex To Concentrate ERK7 in the Apical Cap. mBio: e0286421.

30. Hu, K., D.S. Roos, and J.M. Murray, 2002. A novel polymer of tubulin forms the conoid of Toxoplasma gondii. J Cell Biol, 156: 1039–50.

31. Nichols, B.A. and M.L. Chiappino, 1987. Cytoskeleton of Toxoplasma gondii. J Protozool, 34: 217–26.

32. Nishi, M., K. Hu, J.M. Murray, and D.S. Roos, 2008. Organellar dynamics during the cell cycle of Toxoplasma gondii. J Cell Sci, 121: 1559–1568.

33. Gubbels, M.J., S. Vaishnava, N. Boot, J.F. Dubremetz, and B. Striepen, 2006. A MORN-repeat protein is a dynamic component of the Toxoplasma gondii cell division apparatus. J Cell Sci, 119: 2236–45.

34. Hu, K., 2008. Organizational changes of the daughter basal complex during the parasite replication of Toxoplasma gondii. PLoS Pathog, 4: e10.

35. Frenal, K., D. Jacot, P.M. Hammoudi, A. Graindorge, B. Maco, and D. Soldati-Favre, 2017. Myosin-dependent cell-cell communication controls synchronicity of division in acute and chronic stages of Toxoplasma gondii. Nat Commun, 8: 15710.

36. Engelberg, K., F.D. Ivey, A. Lin, M. Kono, A. Lorestani, D. Faugno-Fusci, T.W. Gilberger, M. White, and M.-J. Gubbels, 2016. A MORN1-assocatiated HAD phosphatase in the basal complex is essential for Toxoplasma gondii daughter budding. Cell Microbiol, 18: 1153–1171.

37. Morano, A.A. and J.D. Dvorin, 2021. The Ringleaders: Understanding the Apicomplexan Basal Complex Through Comparison to Established Contractile Ring Systems. Front Cell Infect Microbiol, 11: 656976.

38. Ouologuem, D.T. and D.S. Roos, 2014. Dynamics of the Toxoplasma gondii inner membrane complex. J Cell Sci, 127: 3320–30.

39. Sheffield, H.G. and M.L. Melton, 1968. The fine structure and reproduction of Toxoplasma gondii. J Parasitol, 54: 209–26.

40. Chiappino, M.L., B.A. Nichols, and G.R. O’Connor, 1984. Scanning electron microscopy of Toxoplasma gondii: parasite torsion and host-cell responses during invasion. J Protozool, 31: 288–92.

41. Morrissette, N., 2015. Targeting Toxoplasma tubules: tubulin, microtubules, and associated proteins in a human pathogen. Eukaryot Cell, 14: 2–12.

42. Leung, J.M., Y. He, F. Zhang, Y.C. Hwang, E. Nagayasu, J. Liu, J.M. Murray, and K. Hu, 2017. Stability and function of a putative microtubule-organizing center in the human parasite Toxoplasma gondii. Mol Biol Cell, 28: 1361–1378.

43. de Leon, J.C., N. Scheumann, W. Beatty, J.R. Beck, J.Q. Tran, C. Yau, P.J. Bradley, K. Gull, B. Wickstead, and N.S. Morrissette, 2013. A SAS-6-like protein suggests that the Toxoplasma conoid complex evolved from flagellar components. Eukaryot Cell, 12: 1009–19.

44. Portman, N. and J. Slapeta, 2014. The flagellar contribution to the apical complex: a new tool for the eukaryotic Swiss Army knife? Trends Parasitol, 30: 58–64.

45. Katris, N.J., G.G. van Dooren, P.J. McMillan, E. Hanssen, L. Tilley, and R.F. Waller, 2014. The apical complex provides a regulated gateway for secretion of invasion factors in Toxoplasma. PLoS Pathog, 10: e1004074.

46. Back, P.S., W.J. O’Shaughnessy, A.S. Moon, P.S. Dewangan, X. Hu, J. Sha, J.A. Wohlschlegel, P.J. Bradley, and M.L. Reese, 2020. Ancient MAPK ERK7 is regulated by an unusual inhibitory scaffold required for Toxoplasma apical complex biogenesis. Proc Natl Acad Sci U S A, 117: 12164–12173.

47. Tengganu, I.F., L.F. Arias Padilla, J. Munera Lopez, J. Liu, P.T. Brown, J.M. Murray, and K. Hu, 2023. The cortical microtubules of Toxoplasma gondii underlie the helicity of parasite movement. J Cell Sci, 136.

48. Tran, J.Q., C. Li, A. Chyan, L. Chung, and N.S. Morrissette, 2012. SPM1 stabilizes subpellicular microtubules in Toxoplasma gondii. Eukaryot Cell, 11: 206–16.

49. Liu, J., Y. He, I. Benmerzouga, W.J. Sullivan, Jr., N.S. Morrissette, J.M. Murray, and K. Hu, 2015. An ensemble of specifically targeted proteins stabilizes cortical microtubules in the human parasite Toxoplasma gondii. Mol Biol Cell.

50. Liu, J., L. Wetzel, Y. Zhang, E. Nagayasu, S. Ems-McClung, L. Florens, and K. Hu, 2013. Novel thioredoxin-like proteins are components of a protein complex coating the cortical microtubules of Toxoplasma gondii. Eukaryot Cell, 12: 1588–99.

51. Liu, P., E. Zupa, A. Neuner, A. Bohler, J. Loerke, D. Flemming, T. Ruppert, T. Rudack, C. Peter, C. Spahn, O.J. Gruss, S. Pfeffer, and E. Schiebel, 2020. Insights into the assembly and activation of the microtubule nucleator gamma-TuRC. Nature, 578: 467–471.

52. Morlon-Guyot, J., M.E. Francia, J.F. Dubremetz, and W. Daher, 2017. Towards a molecular architecture of the centrosome in Toxoplasma gondii. Cytoskeleton (Hoboken), 74: 55–71.

53. M’Saad, O. and J. Bewersdorf, 2020. Light microscopy of proteins in their ultrastructural context. Nat Commun, 11: 3850.

54. M’Saad, O., R. Kasula, I. Kondratiuk, P. Kidd, H. Falahati, J.E. Gentile, R.F. Niescier, K. Watters, R.C. Sterner, S. Lee, X. Liu, P.D. Camilli, J.E. Rothman, A.J. Koleske, T. Biederer, and J. Bewersdorf, 2022. All-optical visualization of specific molecules in the ultrastructural context of brain tissue. bioRxiv: 2022.04.04.486901.

55. Ferreira, J.L., V. Prazak, D. Vasishtan, M. Siggel, F. Hentzschel, A.M. Binder, E. Pietsch, J. Kosinski, F. Frischknecht, T.W. Gilberger, and K. Grunewald, 2023. Variable microtubule architecture in the malaria parasite. Nat Commun, 14: 1216.

56. Padilla, L.F.A., J.M. Murray, and K. Hu, 2024. The initiation and early development of the tubulin-containing cytoskeleton in the human parasite Toxoplasma gondii. Mol Biol Cell: mbcE23110418.

57. Li, Z., W. Du, J. Yang, D.H. Lai, Z.R. Lun, and Q. Guo, 2023. Cryo-Electron Tomography of Toxoplasma gondii Indicates That the Conoid Fiber May Be Derived from Microtubules. Adv Sci (Weinh), 10: e2206595.

58. Heaslip, A.T., F. Dzierszinski, B. Stein, and K. Hu, 2010. TgMORN1 Is a Key Organizer for the Basal Complex of Toxoplasma gondii. PLoS Pathog, 6: e1000754.

59. Lorestani, A., L. Sheiner, K. Yang, S.D. Robertson, N. Sahoo, C.F. Brooks, D.J. Ferguson, B. Striepen, and M.J. Gubbels, 2010. A Toxoplasma MORN1 Null Mutant Undergoes Repeated Divisions but Is Defective in Basal Assembly, Apicoplast Division and Cytokinesis. PLoS ONE, 5: e12302.

60. Stokkermans, T.J., J.D. Schwartzman, K. Keenan, N.S. Morrissette, L.G. Tilney, and D.S. Roos, 1996. Inhibition of Toxoplasma gondii replication by dinitroaniline herbicides. Exp Parasitol, 84: 355–70.

61. Morrissette, N.S. and L.D. Sibley, 2002. Disruption of microtubules uncouples budding and nuclear division in Toxoplasma gondii. J Cell Sci, 115: 1017–25.

62. Reed, S.M., M. Furr, D.K. Howe, A.L. Johnson, R.J. MacKay, J.K. Morrow, N. Pusterla, and S. Witonsky, 2016. Equine Protozoal Myeloencephalitis: An Updated Consensus Statement with a Focus on Parasite Biology, Diagnosis, Treatment, and Prevention. J Vet Intern Med, 30: 491–502.

63. Dubey, R., B. Harrison, S. Dangoudoubiyam, G. Bandini, K. Cheng, A. Kosber, C. Agop-Nersesian, D.K. Howe, J. Samuelson, D.J.P. Ferguson, and M.J. Gubbels, 2017. Differential Roles for Inner Membrane Complex Proteins across Toxoplasma gondii and Sarcocystis neurona Development. mSphere, 2: 00409–17.

64. Brown, K.M., S. Long, and L.D. Sibley, 2018. Conditional Knockdown of Proteins Using Auxin-inducible Degron (AID) Fusions in Toxoplasma gondii. Bio Protoc, 8.

65. Choi, C.P., A.S. Moon, P.S. Back, Y. Jami-Alahmadi, A.A. Vashisht, J.A. Wohlschlegel, and P.J. Bradley, 2019. A photoactivatable crosslinking system reveals protein interactions in the Toxoplasma gondii inner membrane complex. PLoS Biol, 17: e3000475.

66. Bertiaux, E., A.C. Balestra, L. Bournonville, V. Louvel, B. Maco, D. Soldati-Favre, M. Brochet, P. Guichard, and V. Hamel, 2021. Expansion microscopy provides new insights into the cytoskeleton of malaria parasites including the conservation of a conoid. PLoS Biol, 19: e3001020.

67. Beck, J.R., I.A. Rodriguez-Fernandez, J. Cruz de Leon, M.H. Huynh, V.B. Carruthers, N.S. Morrissette, and P.J. Bradley, 2010. A novel family of Toxoplasma IMC proteins displays a hierarchical organization and functions in coordinating parasite division. PLoS Pathog, 6: e1001094.

68. Gajria, B., A. Bahl, J. Brestelli, J. Dommer, S. Fischer, X. Gao, M. Heiges, J. Iodice, J.C. Kissinger, A.J. Mackey, D.F. Pinney, D.S. Roos, C.J. Stoeckert, Jr., H. Wang, and B.P. Brunk, 2008. ToxoDB: an integrated Toxoplasma gondii database resource. Nucleic Acids Res, 36: D553–6.

69. Hawkins, L.M., C. Wang, D. Chaput, M. Batra, C. Marsilia, D. Awshah, and E.S. Suvorova, 2024. The Crk4-Cyc4 complex regulates G(2)/M transition in Toxoplasma gondii. EMBO J.

70. Ferguson, D.J.P. and J.F. Dubremetz, The ultrastructure of Toxoplasma gondii, in Toxoplasma gondii: The Model Apicomplexan, L.M. Weiss and K. Kim, Editors. 2020, Elsevier Academic Press. 21–61.

71. Kohler, S., C.F. Delwiche, P.W. Denny, L.G. Tilney, P. Webster, R.J. Wilson, J.D. Palmer, and D.S. Roos, 1997. A plastid of probable green algal origin in Apicomplexan parasites. Science, 275: 1485–9.

72. Roos, D.S., R.G. Donald, N.S. Morrissette, and A.L. Moulton, 1994. Molecular tools for genetic dissection of the protozoan parasite Toxoplasma gondii. Methods Cell Biol, 45: 27–63.

73. Howe, D.K., M. Yeargan, L. Simpson, and S. Dangoudoubiyam, 2018. Molecular Genetic Manipulation of Sarcocystis neurona. Curr Protoc Microbiol, 48: 20D 2 1–20D 2 14.

74. Granstrom, D.E., O. Alvarez, Jr., J.P. Dubey, P.F. Comer, and N.M. Williams, 1992. Equine protozoal myelitis in Panamanian horses and isolation of Sarcocystis neurona. J Parasitol, 78: 909–12.

75. Louvel, V., R. Haase, O. Mercey, M.H. Laporte, T. Eloy, E. Baudrier, D. Fortun, D. Soldati-Favre, V. Hamel, and P. Guichard, 2023. iU-ExM: nanoscopy of organelles and tissues with iterative ultrastructure expansion microscopy. Nat Commun, 14: 7893.

76. Schindelin, J., I. Arganda-Carreras, E. Frise, V. Kaynig, M. Longair, T. Pietzsch, S. Preibisch, C. Rueden, S. Saalfeld, B. Schmid, J.Y. Tinevez, D.J. White, V. Hartenstein, K. Eliceiri, P. Tomancak, and A. Cardona, 2012. Fiji: an open-source platform for biological-image analysis. Nat Methods, 9: 676–82.

77. Ferguson, D.J., M.F. Cesbron-Delauw, J.F. Dubremetz, L.D. Sibley, K.A. Joiner, and S. Wright, 1999. The expression and distribution of dense granule proteins in the enteric (Coccidian) forms of Toxoplasma gondii in the small intestine of the cat. Exp Parasitol, 91: 203–11.

